# Integrating 3D Tumor Models and Microfluidics for Precise Metabolic Control

**DOI:** 10.64898/2026.01.07.698130

**Authors:** Estelle Bastien, Alpha Diallo, Matthieu Mercury, Jean Capello, Hélène Delanoë-Ayari, Charlotte Rivière

## Abstract

Understanding how metabolic deprivation shapes tumor behavior requires in vitro systems that faithfully reproduce millimeter-scale biochemical gradients. Here, we introduce MilliFlow3D, an open hydrogel-based microfluidic platform that generates controlled metabolic gradients and supports in situ spheroid formation, real-time imaging, and intact retrieval for spatial analyses. Gradients are generated by passive diffusion across a structured agarose microwell array positioned between two perfusion channels. Using a fluorescent tracer and numerical simulations, we show that MilliFlow3D establishes stable linear gradients with local metabolite levels matching those reported in avascular tumor regions. Using HCT116 colorectal cancer spheroids, we demonstrate that a L-glutamine gradient imposes graded effects on growth and proliferation: spheroid expansion decreases from high- to low-glutamine regions, and Ki-67–positive cells progressively shift toward the periphery under glutamine depletion. Thanks to the platform’s spatial accessibility, these phenotypic responses could in the future be coupled to molecular readouts, enabling spatial mapping of metabolic pathway activity along the gradient. Altogether, MilliFlow3D provides a robust and versatile platform to investigate how heterogeneous metabolic landscapes sculpt tumor behavior and to identify context-dependent metabolic vulnerabilities with high analytical precision.

## 1. Introduction

The tumor microenvironment (TME) is a complex and dynamic ecosystem composed of diverse cellular and acellular components. Over the past decade, the TME has been recognized as a key driver of tumor progression and therapeutic resistance, positioning it as a critical target for next-generation cancer therapies. Biochemical hallmarks such as hypoxia, acidosis, and nutrient depletion arise from the dynamic interplay between cancer cells and their microenvironment, and are key drivers of the spatial genetic and phenotypic heterogeneity observed within solid tumors^1,2,3^. Among key metabolites, L-glutamine stands out as the most abundant amino acid in plasma and muscle and a major fuel for cancer cells, making many tumors - including colon tumors - highly dependent, or even “addicted,” to glutamine to sustain proliferation and survival ^4,5^. This strong dependency is accompanied by an upregulation of the glutamine metabolic pathway, notably through increased expression of the ASCT2 transporter ^4^. A key obstacle is the rapid metabolic plasticity of cancer cells, which upregulate glutamine transporters or switch to alternative nutrient sources, ultimately driving aggressiveness and therapy resistance ^6^. These adaptive mechanisms occur within a profoundly glutamine-depleted and highly heterogeneous microenvironment, where L-glutamine levels are markedly reduced in tumors ^7–9^ and unevenly distributed between core and periphery due to poor vascularization and high metabolic demand ^10,11^. Indeed, the abnormal vascularisation within solid tumors will lead to inefficient blood supply, generating nutrient-poor regions and metabolic gradients extending over the millimeter scale ^12,13^.

To investigate how such metabolic deprivation shapes tumor behavior in space and time, we need in vitro systems that can reproduce metabolic landscapes with sufficient analytical detail. Recapitulating these complex metabolic conditions *in vitro* remains challenging. Three-dimensional models such as spheroids and patient-derived organoids improve physiological relevance but remain limited in size (100-500 µm) and therefore cannot reproduce millimeter-scale biochemical gradients characteristic of solid tumors. Development of microfluidic or microphysiological systems are increasing this last decades to address this limitation^14^. They provide precise and reproducible control of heterogeneous biochemical environments, enabling the generation of externally imposed or cell metabolism-driven gradients at tissue-relevant scales^15^. However despite their promise, current microphysiological approaches face several limitations. For instance, the polymer most commonly used, polydimethylsiloxane (PDMS), exibits a strong absorption of therapeutic compounds^16,17^, which undermines reliable drug screening under physiological conditions^18^. Moreover, its impermeability to small water-soluble molecules leads to rapid medium conditioning unless continuous perfusion is maintained. To go beyond PDMS and its limitations, hydrogel-based microwells have been considered ^19,20^, as well as hydrogel-based microphysiological systems^21,22^. Hydrogels are networks of cross-linked polymers with tunable physical properties, a high capacity for water retention, and interconnected pores enabling free diffusion of O_2_, nutrients, and metabolic wastes, which makes them favorable alternatives in micro-system applications. Nevertheless, existing hydrogel-based platforms ^23,24^ are so far relying on injection of cells or pre-formed 3D models through ports on closed system, which (1) prevents accurate positioning within the defined metabolic gradient, (2) complicates sample recovery for downstream analyses, and (3) requires time-consuming construction, meticulous human manipulations and expensive specialized equipment, limiting their use in clinical settings.

To overcome these limitations, we developed an open microfluidic platform "MilliFlow3D" that enables the controlled exposure of 3D tumor models to millimeter-scale gradients of metabolites (see Figure 1). Gradients are generated by diffusion across a structured agarose hydrogel containing a microwell microarray, which enables the direct initiation of spheroids and ensures their precise spatial positioning along the gradient. The open architecture not only facilitates spheroid formation and real-time imaging, but also enables straightforward and intact retrieval of individual 3D models for whole-mount immunostaining, confocal microscopy, and molecular analyses. Agarose was selected for its excellent biocompatibility and the functional tunability it brings to the system ^25,26^. We validated this microphysiological platform by generating controlled millimeter-scale gradients of L-glutamine and assessing their impact on colorectal cancer spheroids, demonstrating its ability to capture spatially resolved metabolic responses with high analytical precision.

**Figure 1.**
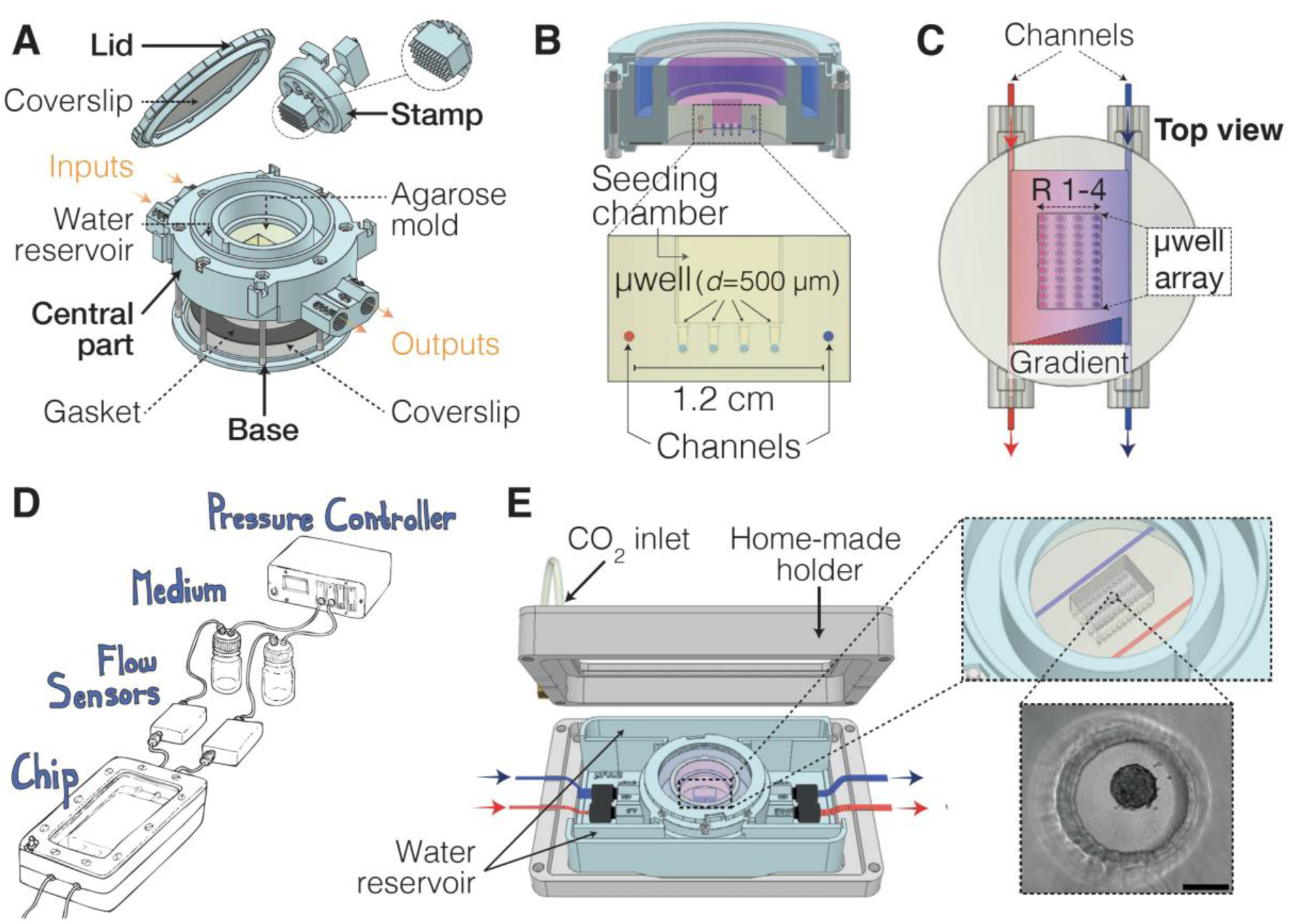
Schematic representation of MilliFlow3D-Chip system. **(A)** Assembly of the MilliFlow3D chip. The system consists of three parts: a central section containing the agarose gel, a peripheral water reservoir, and four luer-lock connectors for inlet and outlet perfusion; a base that seals the device with a coverslip and gasket; and a lid holding a 35-mm glass coverslip (Knittel Glass) enclosed between two printed rings to ensure sterility. A cuboid-shaped stamp with four rows of eleven rods is inserted into the assembled chip, while two stainless-steel needles are positioned between the inlets and outlets. Then, 2% agarose is poured into the central chamber to mold the seeding region containing round micro-wells (500 µm in diameter) and the two parallel perfusion channels. **(B)** Cross-section of the microstructured agarose gel showing two parallel perfusion channels (1 and 2) flanking the central seeding chamber with micro-wells for 3D cell culture. **(C)** Top view of the molded agarose gel integrated within the MilliFlow3D chip. The seeding chamber contains a micro-well array organized in four rows (R1-4) of eleven micro-wells. Perfusion of distinct solutions through channels 1 and 2 generates a stable gradient across the chamber by passive diffusion. **(D)** The chip is connected to a pressure controller via external tubing and **(E)** mounted on a custom holder that maintains controlled CO₂ and humidity conditions through a CO₂ inlet and water reservoir, allowing the monitoring of spheroid growth into microwell (scale bar: 200 μm).

## 2. Results

### 2.1. Design of MilliFlow3D

MilliFlow3D, designed to mimic the scale of tumor tissue, enables the in situ formation of avascular 3D tumor models, their culture under controlled metabolic conditions, and their subsequent analysis (Figure 1A). Unlike closed microfluidic systems, MilliFlow3D features a transparent, unsealed top that preserves sterility while providing direct access to the chip, thereby enabling simple spheroid initiation and easy retrieval for subsequent analyses (Figure 1A–B, Figure 4). The device was fabricated using 3D printing technology. It includes a central part featuring two inlets and two outlets, each fitted with a female Luer-lock connector identical to standard commercial ones, ensuring compatibility with common laboratory setups. The central part, designed to hold the sample of interest, is assembled on a transparent base that hermetically seals the bottom of the chamber. Once assembled, the sealed central chamber serves as a mold for casting the microstructured agarose hydrogel. During the molding step, a custom-designed stamp is inserted into the central cavity, while two stainless-steel needles (outer diameter of 0.6 mm) are positioned between the inlets and outlets to define the parallel perfusion channels (Figure S1A). The stamp consists of a rectangular support bearing rounded-tip microneedles that imprint an array of 500-µm micro-wells (SI Video, Figure S1B-C). After agarose gelation, both the stamp and the needles are removed, resulting in a microstructured agarose gel with a central rectangular region - termed the seeding chamber - containing four rows of eleven micro-wells, with a center-to-center distance of 2 mm between adjacent rows, positioned between the two perfusion channels (Figure 1B, C). These micro-wells enable the in situ initiation and culture of 3D cell models by direct cell seeding into the micro-wells, where cells sediment and self-organize into spheroids, with the culture medium covering the agarose gel. Each channel allows continuous perfusion of the culture medium and maintains a stable concentration of nutrients on both sides of the gel while enabling continuous waste evacuation: one channel is supplied with nutrient-depleted medium, while the other receives nutrient-enriched medium. This controlled concentration difference induces a steady diffusion of solutes across the gel, thereby establishing a stable gradient that enables the culture of spheroids within the micro-well array under spatially defined nutrient conditions.

To minimize agarose drying during long-term experiments, the gel is bordered by a peripheral water reservoir (Figure 1A-B). Additionaly, MilliFlow3D was co-developed with a dedicated microscopy holder that can be fully opened, facilitating sterile connection of the chip to the perfusion setup (Figure 1 D and E).

### 2.2. Establishment and characterization of a stable linear gradient

To validate the ability of MilliFlow3D to establish and maintain a stable gradient across the gel, experiments were performed in empty micro-wells (i.e., without spheroids), we monitored the diffusion of a small soluble molecule, sulforhodamine B (SRB), used as a nutrient-like fluorescent tracer (Figure 2 and Table 1). As illustrated in Figure 2 A-B, the chip was perfused at a flow rate of 4 µL/min with culture medium supplemented with SRB in channel 1 (C_1_), while channel 2 (C_2_) received SRB-free medium. Flow rates were imposed and monitored using a pressure-based control system (ElveFlow) coupled to inline flowmeters. The two channels are 1.2 cm apart and the microwell array (0.7 cm wide) is positioned in the centre. This geometry was specifically designed to generate and resolve gradients over millimetre-scale distances. Live epifluorescence microscopy revealed a progressive increase in fluorescence intensity across the central chamber as time increases, indicating the formation of a linear concentration gradient between the two channels over time. Quantitative analysis of fluorescence profiles demonstrated that the gradient stabilization - defined as a variation below 15% of the final intensity in each microwell row – takes 29 hours of continuous perfusion, as shown by the plateau of both the fluorescence slope and the mean intensity measured at the spatial locations of the four micro-well rows (Figure 2 C and D). Importantly, the uniform and precisely defined spatial organization of the micro-well rows (from rows 1 near C_1_ perfused with SRB to row 4 near C_2_ perfused without SRB) enables reliable quantification of fluorescence along the gradient axis (Figure 2 D). The progressive decrease in fluorescence intensity from row 1 to 4 demonstrates significant differences in SRB concentration between rows, indicating that the micro-well spacing and channel geometry enable the formation of a stable and quantifiable concentration gradient across the micro-well array. Finally, no fluorescence gradient was observed along the flow direction (Figure S2), ensuring homogeneous conditions across each column of the micro-well array and enabling statistically robust repetition within a single chip (n = 11 micro-wells per row).

**Figure 2.**
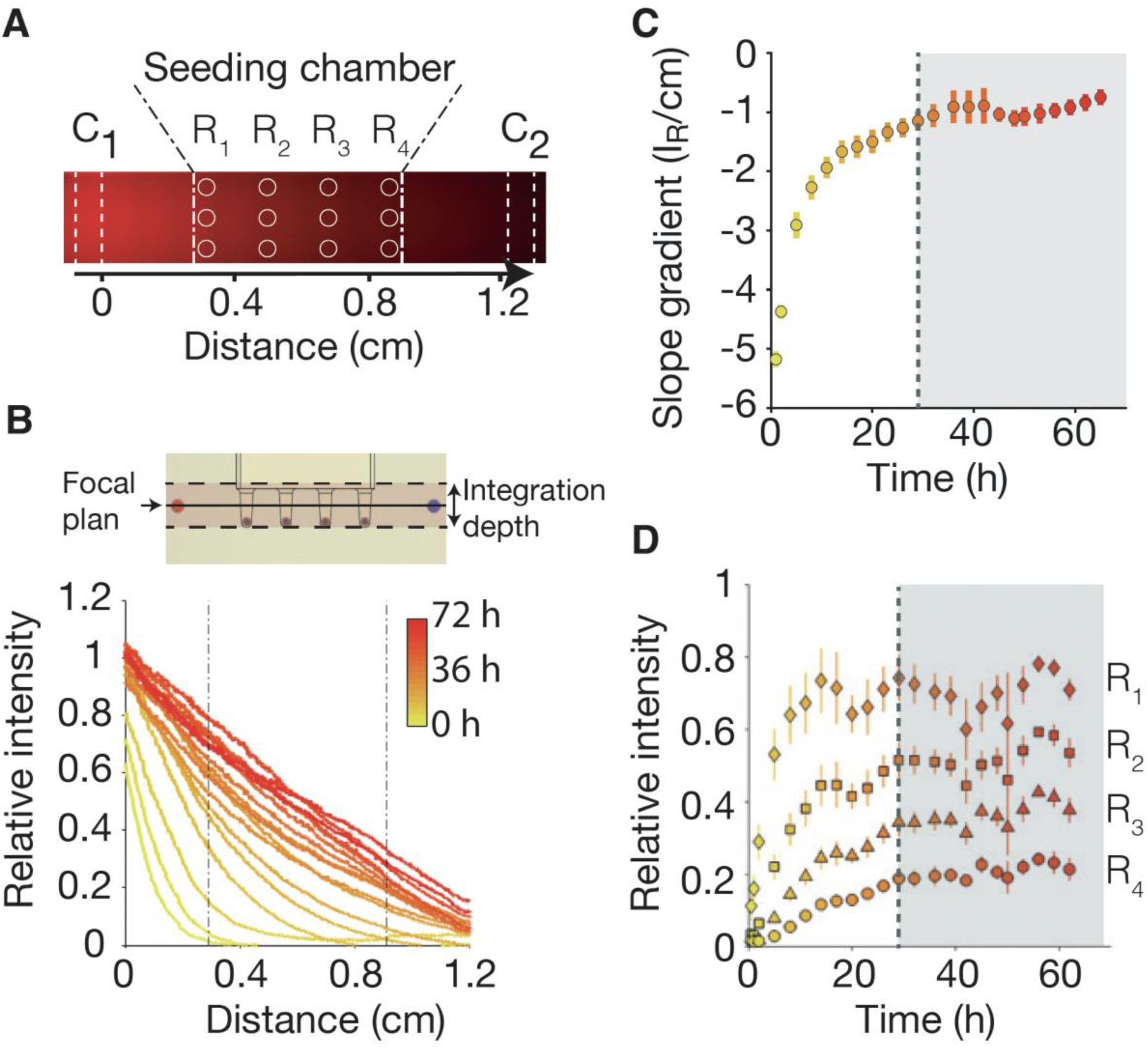
Stable linear gradient of fluorescent tracer. **(A)** Stitched epifluorescence images of the central region of the chip acquired from the bottom of the chip, showing the perfusion channels at the image edges, the seeding chamber at the center (outlined with dashed lines), and the locations of the microwells (circles; rows R1–R4). The chip was perfused for 72 h at 4 µL.min⁻¹ per channel with SRB (1 µg.mL⁻¹) in channel 1 and dye-free medium in channel 2. **(B)** SRB fluorescence intensity profiles between channels 1 and 2, recorded every 3 h (0-72 h, color gradient yellow to red). Dashed lines indicate the location of the seeding chamber. Top: schematic illustration of the focal plane positioned at the channel center and of the 2-mm fluorescence signal integration depth. (C) Temporal evolution of the profile slope quantified between 0.25 and 0.35 cm from channel 1. Mean +/- SEM (D) Time course of the mean fluorescence intensity averaged over a 500-µm region at the microwell rows (R1-R4). Mean +/-SEM. In (C) and (D), the vertical grey line indicates the gradient stabilization point (grey area). (*N* = 4).

**Table 1.** Parameters used for gradients establishement simulation.

| Parameter | Abbreviation | Value | Reference |
| --- | --- | --- | --- |
| Agarose 2% permeability | $k$ | $1 \times 10^{-10} \text{ m}^2/\text{s}$ | 25 |
| Agarose 2% porosity | $\phi$ | 0.905 | 25 |
| Channel diameter | $d$ | 0.6 mm | |
| Flow rate | | 4 $\mu\text{l}/\text{min}$ | a |
| SRB diffusivity | $D_{SRB}$ | $1.06 \times 10^{-9} \text{ m}^2/\text{s}$ | 28 |
| Glutamine diffusivity | $D_{Glut}$ | $1.02 \times 10^{-9} \text{ m}^2/\text{s}$ | 29 |
| Glutamine cell consumption | $q$ | $1.6 \times 10^{-2} \text{ fmol}/\text{cell}/\text{h}$ | 30 |
| Cell number/spheroid | $n$ | 4000 | a |
| Spheroid diameter | | 400 $\mu\text{m}$ | |
| Spheroid consumption | | 1.9 mmol/$\text{m}^3/\text{s}$ | |

<sup>a</sup> Experimental Data

### 2.3 Numerical modeling of L-glutamine gradient formation within MilliFlow3D

While small-molecule diffusion could be experimentally validated using SRB, direct visualization of nutrient gradients such as L-glutamine remains challenging. To address this limitation, we performed numerical simulations using COMSOL Multiphysics to model gradient establishment within MilliFlow3D, taking into account both physical parameters - including gel properties, diffusion coefficient, and initial L-glutamine concentration - and biological aspects such as spheroid cell density and glutamine consumption rate (Table 1, Figure 3 A). The standard DMEM medium used for HCT116 culture and spheroid initiation in the chip contains 2 mM L-glutamine, saturating the agarose matrix at 2 mM (Figure 3A, C₂, burgundy) prior to perfusion. To mimic physiologically relevant conditions (0.7–0.2 mM *in vivo*^8,27^), the medium covering the gel was replaced with glutamine-depleted medium (Figure 3 A, C₀, dark blue) to reduce the residual glutamine reservoir within the chip, and the two channels were perfused with media containing 1 mM (C₁, green) and 0 mM (C₀) glutamine, respectively (Figure 3 A). Because cells actively consume glutamine, a maximal homogeneous consumption rate per spheroid (Table 1) was applied to test whether local uptake would alter gradient stability.

**Figure 3.**
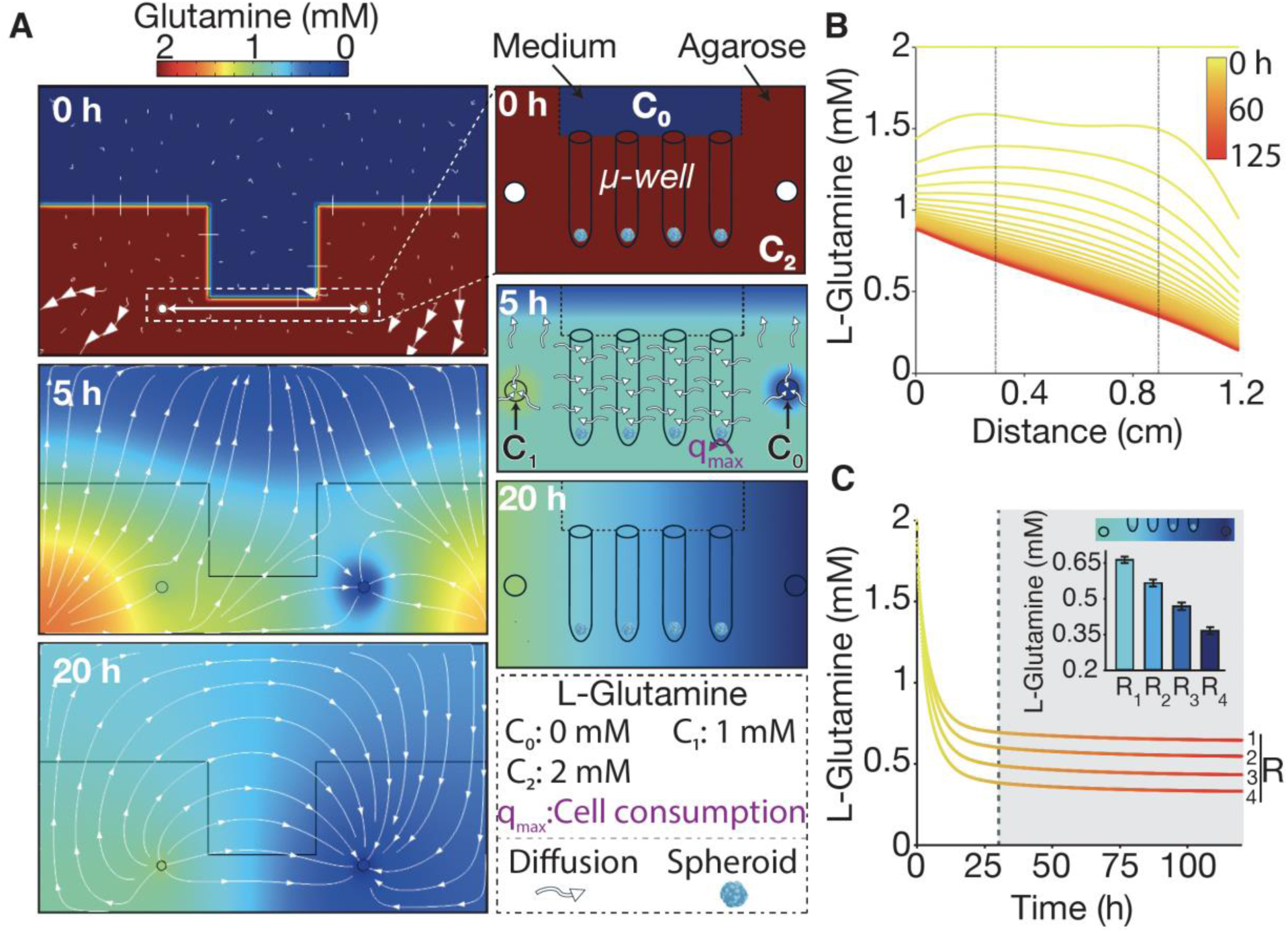
Simulation of L-glutamine gradient establishement. (**A**, left) 2D diffusion-simulated cross-sectional views showing the establishment of the L-glutamine gradient over time within the agarose gel containing the seeding chamber and the two perfusion channels (1 and 2) supplied with 1 and 0 mM L-glutamine, respectively. The color map represents L-glutamine concentration, and arrows indicate diffusion fluxes. (**A**, right) Schematic representation of the simulation parameters. The model geometry comprises four structural components: (1) the agarose layer containing four microwell rows within the seeding chamber, overlaid by (2) the medium layer, and (3–4) the two perfusion channels. The color map and arrows indicate L-glutamine concentration and its diffusion gradients, respectively: C_0_ (0 mM) corresponds to the medium overlying the agarose gel and perfusing channel 2; C_1_ (1 mM) to the medium perfusing channel 1; and C_2_ (2 mM) to the initial concentration within the agarose gel. Spheroids embedded in the agarose layer were defined as active consumers with a maximal L-glutamine uptake rate. (**B**) L-glutamine concentration profiles extracted from the 3D simulations between channels 1 and 2 (along the double-arrow line shown in up panel A), recorded every 1 h from 0 to 125 h (color gradient from yellow to red). Dashed lines indicate the position of the seeding chamber. (**C**) Time course of the mean L-glutamine concentration computed from the 3D simulation results, averaged over a 500-µm region at the microwell rows containing spheroids (R1-R4). The vertical grey line indicates the gradient stabilization point (grey area). *Inset*: averaged L-glutamine concentration between 25 and 125 h as a function of microwell row position (R1-R4). Standard deviation represent the difference of concentration experienced by each spheroids in the defined row.

Initial concentration differences between the channels, the agarose matrix, and the covering medium generated diffusion streamlines that progressively stabilized over time (Figure 3A). At 5 hours post-perfusion, the L-glutamine gradient displayed a marked asymmetry, caused by the unequal concentration differentials between each channel and the agarose. The 1 mM L-glutamine in C_1_ produces a smaller concentration difference with the agarose than the 0 mM in C_2_, thereby generating a transient asymmetric profile. A steady and linear L-glutamine gradient formed between the two channels, reaching stabilization - defined as a variation below 15% of the final concentration in each microwell row - after 29.5 hours of perfusion (Figure 3 B and C). Simulated local L-glutamine concentrations ranged from 0.65 +/- 0.02 to 0.35 +/- 0.02 mM across microwell rows 1 to 4, spanning the 0.7 cm distance between the row closest to C₁ and the row closest to C₀ (Figure 3 C). These values fall within the physiological range reported in tumor tissues. Importantly, the linearity and stability of the gradient remained intact even under maximal cellular consumption, confirming the robustness of the MilliFlow3D design.

### 2.3 L-glutamine gradient controlled spheroid growth

The MilliFlow3D chip provides an integrated platform for spheroid generation and controlled L-glutamine gradient exposure (Figure 4). Following cell seeding into the micro-wells, cells sediment and self-organize into spheroids with an average diameter of 248 µm ± 31 µm (Figure 4 A and B). Two days after spheroid formation, the MilliFlow3D chip is connected to the perfusion circuit, leading to the establishment of a stable L-glutamine gradient within 29 hours, as predicted by numerical simulation (Figure 3). Spheroids distributed across rows 1 to 4 along the perfusion axis - ranging from the channel perfused with 1 mM L-glutamine to micro-wells deprived of L-glutamine - are monitored by live video microscopy for four additional days prior to subsequent off-chip analyses. Exposure to the L-glutamine gradient markedly affects spheroid growth (Figure 4 C and D). Growth rates decrease progressively from row 1 to row 4, reflecting the spatially decreasing L-glutamine concentrations. This behavior is qualitatively consistent with the growth pattern observed in ULA plates at L-glutamine concentrations corresponding to the simulated values for each row (Figure S3 A). Importantly, when both channels are perfused with 1 mM L-glutamine, thereby eliminating the gradient, no significant growth differences are observed between the rows (Figure S3 B), confirming that the spatial growth heterogeneity arises from the imposed metabolic gradient rather than from intrinsic positional effects within the chip. This is further supported by a two-dimensional COMSOL simulation demonstrating homogeneous stabilization of L-glutamine at 1 mM throughout the chip under symmetric perfusion conditions (Figure S3 D). After four days of perfusion, the central region of the microfluidic chip - corresponding to the rectangular chamber where spheroids were cultured – was retrieved using a custom-made puncher (Figure 4 E), processed as a whole for immunostaining. This procedure preserves the spatial organization of the four microwell rows, allowing each spheroid to be imaged and analyzed according to its precise row position along the gradient (Figure S4). Reduced growth is accompanied by a spatial restriction of proliferative cells. After four days under the L-glutamine gradient, Ki-67 staining indicates that proliferative activity becomes confined to the spheroid periphery (Figure 4 F-I). Ki-67 signal within the core decreases progressively from row 1 to row 4, in parallel with the decreasing metabolic availability (Figure 4 I).

**Figure 4.**
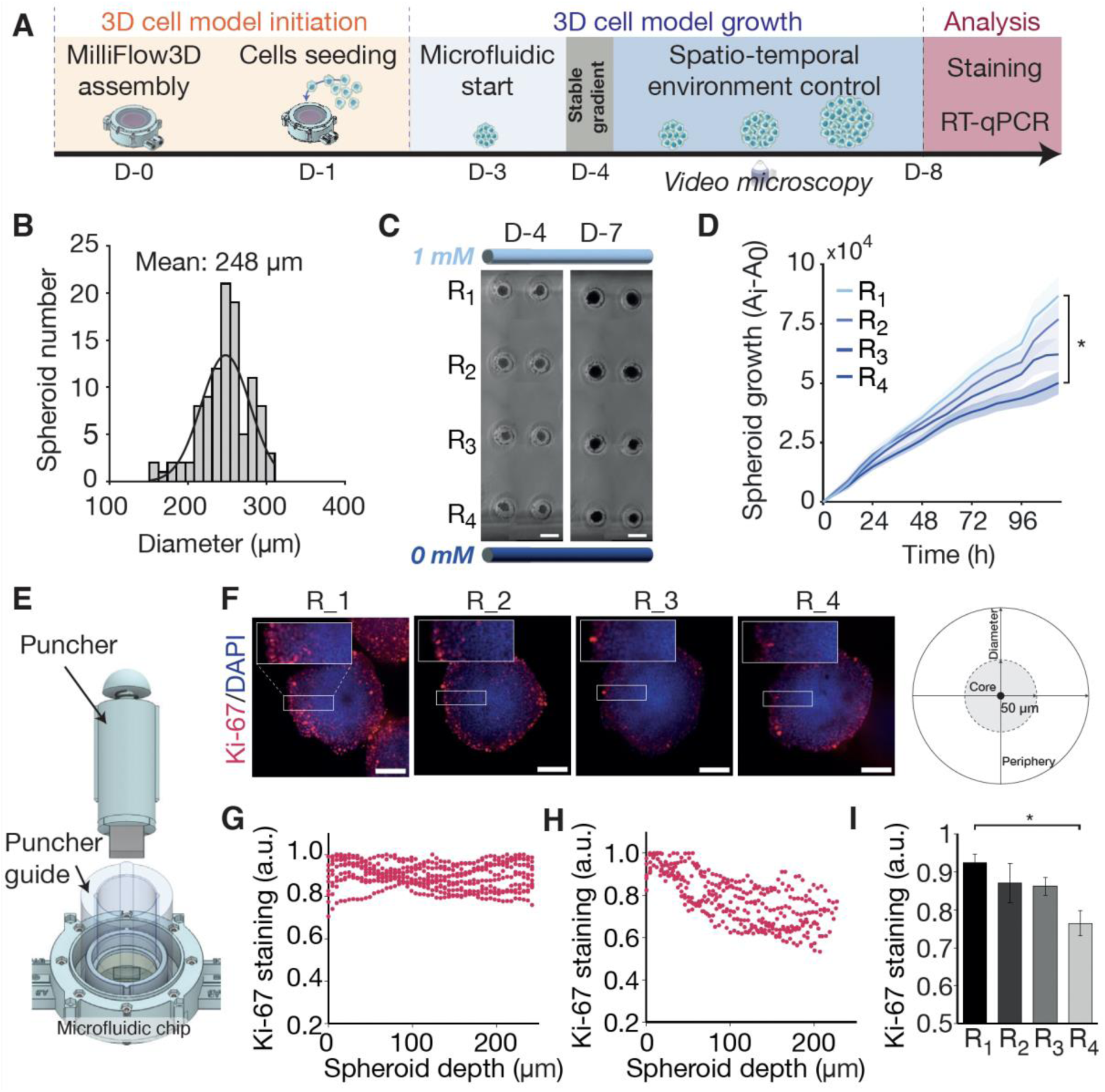
Spheroid growth in the MilliFlow3D chip under L-glutamine gradient. (**A**) Experimental timeline: chip assembly and spheroid seeding, followed by a 3-day static culture. Microfluidic perfusion was then initiated to establish an L-glutamine gradient within 24 h, and spheroid growth was monitored until day 8, when spheroids were collected for IF or RT-qPCR analyses. (**B**) Distribution and mean diameter of 3-day-old HCT116 spheroids quantified from brightfield images using Fiji (*N* = 11, *n*_spheroid_= 44). (**C**) Representative stiching of brightfield images of two HCT116 spheroids per row, acquired at gradient stabilization (day 4) and three days later (day 7), scale bar: 500 μm. (**D**) Normalised spheroid growth (A-A_0,_ where A is the spheroid area at time *t* and A_0_ its initial area) as a function of their position in the chip (rows 1 to 4). Data are presented as mean ± SEM (*N* = 4). (E) Custom-made puncher used to extract the central region (seeding chamber) of the agarose gel containing the spheroid rows at the end of the experiment for downstream analyses. (**F**) Ki-67 immunostaining of 8 day-old spheroids cultured in MilliFlow3D (scale bar: 100 μm). (**G-H**) Single-spheroid normalized (I/I_max_) radial profiles of Ki-67 staining from the periphery to the core, on spheroids cultured in row 1 (**G**) et 4 (**H**). (**I**) Normalized mean Ki-67 intensity measured within a 50-µm-diameter core region across rows 1 to 4. Data are presented as mean ± SEM (*N* = 2, *n*_spheroid_>20 per condition).

### 2.4 Spatially resolved sample preparation for molecular analyses

To enable spatially resolved molecular analyses of spheroids cultured under nutrient gradients, we developed a dedicated workflow for sample retrieval and spatial segmentation within the microfluidic chip. After four days of perfusion, the central region of the chip - corresponding to the rectangular chamber containing the spheroids - was retrieved using a custom-made puncher (Figure 4E). The resulting agarose block was placed on a cutting guide and divided into four longitudinal strips, each corresponding to one row of micro-wells. Each strip was then carefully inverted and positioned onto a dedicated support (strip holder; Figure 5A). The assembly was transferred into a 12-well plate, and a brief centrifugation step enabled the recovery of spheroids according to their spatial position along the gradient (rows 1-4). This workflow provides spatially resolved spheroid samples compatible with downstream molecular analyses, including gene-expression profiling, thereby enabling future investigation of nutrient-dependent molecular programs within the same microfluidic platform.

**Figure 5.**
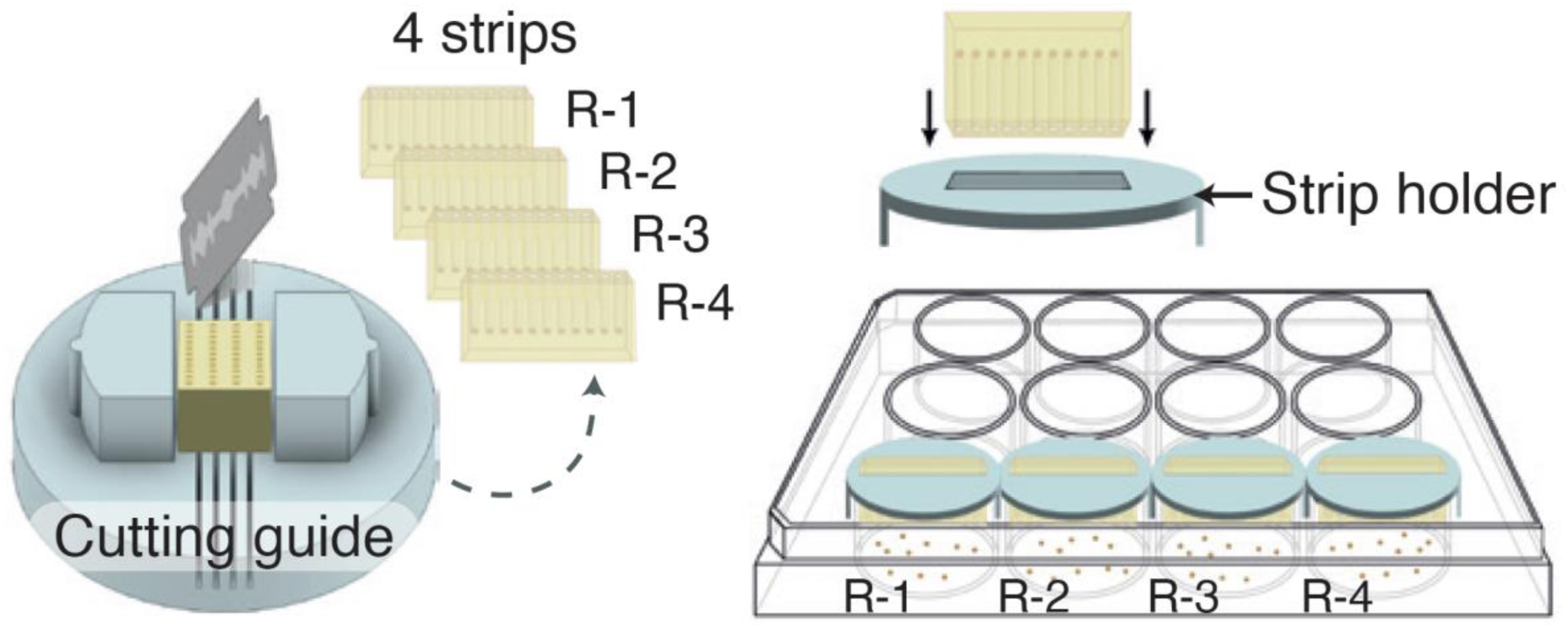
Spatial recovery of spheroids for downstream molecular analyses. (A) Workflow for spatial spheroid isolation. The central agarose block containing the seeding chamber is excised using a custom puncher, then positioned on a cutting guide and sectioned into four longitudinal strips, each corresponding to one microwell row. Each strip is gently inverted onto a dedicated strip holder placed above a 12-well plate. A brief centrifugation releases the spheroids into the underlying wells, enabling their collection according to spatial position for downstream processing such as RT–qPCR.

## 3. Discussion

The MilliFlow3D system presented in this article is an all-in-one platform enabling 3D cell models, as spheroids, to be exposed to controlled metabolic landscapes over several days. A stable linear gradient is generated by passive diffusion between the two perfusion channels, while the linear arrangement of the 3D cultures within the microwell array allows precise correlation between their position and the local metabolite concentration. Unlike conventional PDMS-based or closed hydrogel microfluidic systems, the open architecture of MilliFlow3D provides direct physical access throughout the experiment, enabling straightforward seeding, real-time imaging, and easy retrieval of intact 3D models for downstream immunostaining or molecular analyses. Together, these features enable high-resolution analysis of cellular responses to nutrient levels across the gradient in an avascular tumor context.

Using glutamine-addicted HCT116 spheroids as a test case, we demonstrate that controlled L-glutamine gradients established in MilliFlow3D are sufficient to impose graded effects on tumor growth and on the spatial organization of proliferative cells within the 3D structure. Importantly, the gradient covers physiologically relevant concentrations, mirroring the nutrient-depleted conditions typically found in poorly vascularized tumor regions (0.7–0.2 mM ^8,27^). Spheroid growth progressively decreases from row 1 to row 4, aligning with the growth observed in ULA controls at concentrations corresponding to the simulated local glutamine levels. This controlled spatial response is enabled by the centimeter-scale geometry of the device, which generates a linear gradient from 0.65 +/- 0.02 to 0.35 +/- 0.02 mM over 6 mm. At the scale of individual spheroids (< 500 µm in diameter, matching the 500-µm microwells), the local glutamine concentration varies only minimally - by less than ∼0.03 to 0.05 mM across a spheroid - ensuring that the observed responses arise from global positional differences rather than intra-spheroid fluctuations. Beyond global growth effects, Ki-67 profiling reveals a shift toward peripheral cell proliferation in small spheroids (< 500 µm) under low-glutamine exposure, recapitulating hallmark features of poorly vascularized tumor regions where nutrient deprivation shapes spatial cell-cycle dynamics. Thanks to the spatial accessibility of the MilliFlow3D platform, these phenotypic changes can be directly connected to their underlying molecular programs: the central agarose block can be retrieved, sectioned according to microwell rows, and processed for whole-mount immunostaining or gene-expression analysis, enabling precise spatial mapping of metabolic pathway activity - such as glutamine transport and catabolism - along the gradient, with each row comprising 11 micro-wells that provide statistically robust intra-chip replication.

Such L-glutamine deprivation sensitivity is fully aligned with the metabolic biology of colorectal cancer, which frequently displays strong glutamine dependence^4,5^. As a result, even modest changes in extracellular glutamine availability can impact cell phenotype, making MilliFlow3D particularly suited to dissect how metabolic gradients structure tumor behavior. To go further in order to mimic tumor tissue characteristics, the tunability of agarose mechanical properties offers the unique opportunity to match the stiffness of different tumor tissues. This opens the possibility to systematically vary both matrix properties and metabolic inputs to interrogate how biophysical and biochemical factors jointly regulate tumor behavior. In the same way, adjusting agarose porosity allows us to modulate diffusion behavior and to mimic the transport of free or nanoscale chemotherapeutic agents through dense tumor ECM. Combined with the simulation framework already implemented, MilliFlow3D could ultimately be used to predict diffusion profiles under desired pathological conditions, reducing experimental burden while guiding therapeutic design.

Microfluidics integrated with 3D cell models has led to the emergence of numerous organ-on-chip systems^31^ addressing tumor-stromal interactions^32^, metabolic crosstalk, or vascularization-driven growth^33^. MilliFlow3D complements these approaches by offering a simpler, open, and highly tunable platform specifically optimized for controlled metabolic perturbations at the millimeter scale, an intermediate regime that remains difficult to capture with classical microfluidics or spheroid assays.

Although this study focused on a single colorectal cancer cell line and avascular spheroids, these models provide a relevant starting point to probe metabolic dependencies. The platform is readily compatible with patient-derived organoids, stromal or immune co-cultures, and additional gradients such as oxygen or lactate. Furthermore, replacing agarose with colonizable matrices such as collagen^34^ or GelMA^35^ would allow functionalization of peripheral channels with endothelial cells, enabling the reconstruction of perfusable vascular-like structures more closely reproducing the pathophysiology of solid tumors.

Altogether, MilliFlow3D provides a robust and flexible framework to dissect how spatially heterogeneous metabolic landscapes sculpt tumor growth through metabolic adaptation, and to identify context-dependent metabolic vulnerabilities in a controlled, yet physiologically relevant, setting.

## 4. Conclusion

The development of the MilliFlow3D platform results from an innovative approach combining engineering, computational, and biological techniques synergistically, to offer an enhanced capability for studying tumor metabolism and responses to environmental changes. By integrating microfluidics with 3D cancer spheroid or organoid models that mimic avascular tumors, our system offers a powerful platform for identifying novel metabolic therapeutic targets with unprecedented precision and ease.

## 5. Experimental Section/Methods

### Reagents

Standard agarose, Sulfo-Rhodamine B (SRB), D-glucose and HEPES were purchased from Sigma-Aldrich. L-Glutamine was obtained from Gibco.

### Cell culture

HCT-116 (#CCL-247) a human colon cancer cell line was purchased from American Type Culture Collection (ATCC). Cells were stored according to the supplier’s instructions and used within 6 months after resuscitation of frozen aliquots. Cells were maintained in DMEM-Glutamax (2 mM) media supplemented with 10% of heat-inactivated FBS, and 1% of Pen/Strep. Cells were tested for mycoplasma contamination with MycoAlert Mycoplasma Detection kit (#LT07, Lonza) before being used.

### Microfluidic system setup

The MilliFlow3D Chip design was created using Fusion 360 CAD software, and the chip components were subsequently fabricated using biocompatible High-Temp resin (Formlabs) and Formlabs Form-3 3D printer. This resin was chosen because of its compatibility with autoclave, as well as its biocompatibility tested in different cellular context^36,37^. As shown in **Figure 2 A**, the chips consist of three parts: (1) the central part, (2) a lid, and (3) a base. The central part carries the main compartment where the agarose hydrogel will be cast, surrounding water reservoir to avoid evaporation, and four luer-lock connectors to connect external microfluidic tubing with standard ¼-28 male luer-lock connectors. Two cylindrical stainless-steel rods (0.6 × 80 mm, BRAUD) are inserted through the female luer-lock connectors to mold two parallel channels (600 µm in diameter) within the hydrogel, separated by a 12 mm center-to-center distance. A stamp with a cuboid shape (7×11 mm, **Figure 2 A**), featuring four columns of eleven rods, each 2 mm long and 500 µm in diameter, with rounded tips, is inserted into the central chamber, creating the array of micro-wells necessary for the *in-situ* initiation of 3D cell cultures (Figure S1 B). The stamp design is optimize to match the chip inner agarose chamber, ensuring reproducible positioning of the wells. The diameter and rounded tips of the stamp were characterized using optical profilometry (Contour GT Bruker) with a G=10x objective (**Figure S1 C and D**). In the paper, the cuboid with the micro-wells is referred to as the seeding chamber (**Figure 2 B**). The bottom of the central part is sealed by sequentially adding a silicone flat gasket (0.5 mm thick, Grace Bio-Labs), a coverslip (n°1, thickness 170 µm, diam. 35 mm, Knittel Glass) and the base, all of which are secured together using eight screws.

Once the chip was assembled, hydrogels were molded by dispensing a 2% (w/v) agarose solution pre-warmed to 110 °C into the central compartment. After gelation (25 min at room temperature), the stamp and rods were carefully removed. The microfluidic chip was then filled with PBS and sterilized under UV light (254 nm, 60 W, 30 min). The top of the chip was closed with a lid consisting of a 35 mm glass coverslip (n°1, thickness 170 µm, Knittel Glass) enclosed between two 3D-printed rings. This lid design ensured sterility during cell culture and allowed gas exchange, facilitated by a small stud on the surface in contact with the chip (Figure 2A and Figure S1-A).

Twenty-four hours before the experiment, PBS was replaced with culture medium, and the chips were incubated at 37 °C to allow agarose saturation with the medium. Microfluidic perfusion was driven by a pressure controller (ElveFlow) at a flow rate of 4 μL.min⁻¹ per channel, monitored using flow sensors (ElveFlow) and by hourly weighing of the collected medium from the outlets (Figure S1 E). Each channel was connected to an independent reservoir containing the appropriate culture medium to be perfused through channels 1 and 2 (Figure 2B). The outlet tubing from each channel was immersed in a common waste reservoir, ensuring continuous drainage of the perfused medium. Because the top of the chip is not sealed, the pressure at the gel surface corresponds to atmospheric pressure. Under these conditions, even small hydrostatic height differences between the chip, and the waste reservoir can induce counter-flow rates comparable to or exceeding the imposed perfusion rate. Therefore, particular care was taken to align the vertical position of the agarose chip with that of the waste reservoir. In addition, a large-volume waste reservoir was used to prevent any change in medium height over time. The absence of counter-flow was systematically verified by combining inlet flow-rate monitoring using in-line sensors with outlet flow quantification by gravimetric measurements over time.

For live microscopy, the chip was positioned on a custom-made holder (Figure 2 C) that enabled sterile connection to the perfusion system inside a laminar flow hood through its openable design. The holder also maintained environmental conditions (37 °C, 5% CO₂, high humidity) during imaging, the latter ensured by a built-in water reservoir.

### Gradient characterization

To characterize the establishment of concentration gradients within the agarose hydrogel, Sulfo-Rhodamine B (SRB, 558.7 Da, 1 µg.mL⁻¹) was added to the culture medium perfusing channel 1. Gradient formation was monitored every 30 min over 72 h by epifluorescence microscopy (G = 5×, ON = 0.12, Leica DMi8). The entire microfluidic chamber was reconstructed by stitching individual images based on the recorded stage coordinates, with a minimum overlap of 10%. For overlapping regions, the maximum intensity value was retained. In the reconstructed images, fluorescence intensity profiles were extracted along lines drawn perpendicularly between the two channels. Gradient stability was assessed by (i) measuring the slope of a linear fit applied to the initial linear region of the fluorescence profile starting from channel 1, and (ii) monitoring the temporal evolution of the mean fluorescence intensity averaged over a 500-µm region corresponding to the position of the microwell rows. Image analysis was performed using custom scripts in MATLAB. Gradient stabilization time was defined as the earliest time point at which the normalized SRB fluorescence intensity measured in all four microwell rows fell within 15% of their final steady-state value.

### Spheroid culture under controlled L-glutamine conditions

Spheroids were initiated either in the MilliFlow3D Chip or in cell-repellent 96-well plates (Greiner Bio-One), used as gold-standard controls. For microfluidic cultures, 2.1×10⁵ cells were seeded into the central seeding chamber and allowed to settle into the micro-wells under orbital agitation (250 rpm, 20 min), yielding approximately 500 cells per micro-well. Untrapped cells were removed, 4 mL of DMEM was added, and the chip was centrifuged. After a 2-day incubation, the MilliFlow3D Chip was perfused for five days with phenol red-free DMEM, with channel 1 supplemented with 1 mM L-glutamine and channel 2 containing L-glutamine-free medium. One day after the onset of perfusion, defined as day 0 when the L-glutamine gradient was established, spheroid growth was continuously monitored by brightfield time-lapse microscopy (Leica DMi8, 30 min intervals). In parallel, 500 HCT116 cells per well were seeded in 200 µL DMEM in 96-well ULA plates. Three days after seeding (day 0), spheroids were cultured in phenol red-free DMEM containing decreasing L-glutamine concentrations, with half of the medium renewed every 2–3 days. Growth dynamics were assessed by brightfield microscopy (Leica DMIRB). In both systems, spheroid volume was quantified using automated intensity thresholding in MATLAB.

### Recovery of spheroids for immunostaining

After 4 days of culture under L-glutamine gradient thanks to the microfluidic perfusion, the agarose hydrogel containing the spheroids was recovered using a home-made puncher (design with Fusion 360 CAD software and fabricated using Formlabs Form-2 3D printer; Figure 5). For immunostaining, 3D cell models were fixed with 4% paraformaldehyde, and incubated for 3 days with anti-Ki67 (#55609, BD Biosciences, 1/500) antibody. After 3 washing steps with washing buffer (PBS, 0.1% Triton 100-X) in PBS, spheroids were then incubated for 4 days with secondary antibodies (Invitrogen, 1/250), Alexa Fluor 488-conjugated phalloidin (#A12379, Invitrogen, 1/250) to stain actin, and nucleus were counterstained with DAPI (1/5000). Spheroids were then cleared using Glycerol-Fructose-Urea (FunGi) solution according to published protocol ^38^. Staining was visualized by spinning disk confocal microscopy (Nikon) and quantified in three dimension using MATLAB.

### Numeric Simulation

Diffusion of L-glutamine and SRB in the agarose-based microfluidic system was simulated using COMSOL Multiphysics®, considering parameters listed in Table 1, including the porosity and permeability of 2% agarose hydrogel (Pluen et al., 1999), diffusion coefficients of L-glutamine in agarose, and microfluidic chip geometrical and flow properties (dimensions, channel diameter, and applied pressure). The simulation was run over a period of 72 h with a 1 h time step, starting from the onset of perfusion. Channels 1 and 2 were perfused with media containing 1 mM and 0 mM L-glutamine, respectively. Because cells actively consume glutamine, this biological parameter was incorporated into the model. Cell density was estimated by dissociating spheroids of approximately 400 µm in diameter and counting individual cells. A maximal consumption rate per spheroid was then applied, assuming homogeneous uptake among cells (Table 1, 1.6^10^-2^ fmol/cell/h, ^39^), to test the most stringent condition. Three-dimensional simulations were performed using the *Transport of Diluted Species* module coupled with *Free and Porous Media Flow*, enabling the computation of concentration profiles established between the two perfused channels across the agarose hydrogel. Gradient stabilization time was defined as the earliest time point at which the simulated L-glutamine concentration in all four microwell rows fell within 15% of their final steady-state value. In addition, two-dimensional simulations considering only diffusive transport of L-glutamine were conducted to generate illustrative diffusion maps under symmetric perfusion conditions.

### Statistical analysis

Results are expressed as mean ± SEM of at least three independent experiments (*N*), unless otherwise specified. Sample sizes (*n*) are indicated in the figure legends. Statistical analyses were performed in MATLAB. Data normality was assessed using the Lilliefors test, and group comparisons were evaluated by one-way ANOVA followed by Tukey–Kramer post hoc tests. Significance was set at *p* < 0.05.

## Acknowledgements

The authors would like to acknowledge A. Piednoir for her help in the microwells characterisation. We also acknowledge the contribution of SFR Santé Lyon-Est (UAR3453 CNRS, US7 Inserm, UCBL) facility : CIQLE (a LyMIC member), where confocal acquisition were performed, the µFab platform for microfluidic chip design and printing, as well as the 3D-ONCO platform for fruitfull discussion and advise of tumoroïd model (Institut Convergence Plascan, Centre de Recherche en Cancérologie de Lyon, INSERM U1052-CNRS UMR5286, Centre Léon Bérard). This work was supported by the Institut Convergence Plascan (ANR-17-CONV-0002), the IMITATE project (ANR-22-CE51-0043-01), operated by the French National Research Agency (ANR), and the Institut Universitaire de France (IUF). It was also financially supported by the CNRS MITI (MATISSE Project). The salary of E. Bastien was supported by the University Lyon 1 (ETOILE 2022 funding and PACTE project). J.C. acknowledge support by F.R.S.-FNRS under the research grants CR n◦40017301 “G.El.In.Flow”.

## Bibliographie

1. Jin, M.-Z. & Jin, W.-L. The updated landscape of tumor microenvironment and drug repurposing. Sig Transduct Target Ther 5, 166 (2020).

2. Mbeunkui, F. & Johann, D. J. Cancer and the tumor microenvironment: a review of an essential relationship. Cancer Chemother Pharmacol 63, 571–582 (2009).

3. Carmona-Fontaine, C. et al. Metabolic origins of spatial organization in the tumor microenvironment. Proc. Natl. Acad. Sci. U.S.A. 114, 2934–2939 (2017).

4. Watanabe, T. et al. Differential gene expression signatures between colorectal cancers with and without KRAS mutations: Crosstalk between the KRAS pathway and other signalling pathways. European Journal of Cancer 47, 1946–1954 (2011).

5. Fan, Y. et al. Exploiting the Achilles’ heel of cancer: disrupting glutamine metabolism for effective cancer treatment. Front. Pharmacol. 15, 1345522 (2024).

6. Lv, H. et al. Glutamine: a new strategy for targeted metabolic therapy in the tumor microenvironment. Cell Death Discov. 11, 459 (2025).

7. Roberts, E., Simonsen, D. G., Tanaka, K. K. & Tanaka, T. Free amino acids in growing and regressing ascites cell tumors: host resistance and chemical agents. Cancer Res 16, 970–978 (1956).

8. Rivera, S., Azcón-Bieto, J., López-Soriano, F. J., Miralpeix, M. & Argilés, J. M. Amino acid metabolism in tumour-bearing mice. Biochemical Journal 249, 443–449 (1988).

9. Márquez, J., Sánchez-Jiménez, F., Medina, M. A., Quesada, A. R. & De Castro, I. N. Nitrogen metabolism in tumor bearing mice. Archives of Biochemistry and Biophysics 268, 667–675 (1989).

10. Pan, M. et al. Regional glutamine deficiency in tumours promotes dedifferentiation through inhibition of histone demethylation. Nat Cell Biol 18, 1090–1101 (2016).

11. Reid, M. A. et al. The B55α Subunit of PP2A Drives a p53-Dependent Metabolic Adaptation to Glutamine Deprivation. Molecular Cell 50, 200–211 (2013).

12. Vaupel, P. & RallinoÂ, F. Blood Flow, Oxygen and Nutrient Supply, and Metabolic Microenvironment of Human Tumors: A Review.

13. Guzelsoy, G. et al. Cooperative nutrient scavenging is an evolutionary advantage in cancer. Nature 640, 534–542 (2025).

14. Leung, C. M. et al. A guide to the organ-on-a-chip. Nat Rev Methods Primers 2, 33 (2022).

15. Ayuso, J. M. et al. Tumor-on-a-chip: a microfluidic model to study cell response to environmental gradients. Lab Chip 19, 3461–3471 (2019).

16. van Meer, B. J. et al. Small molecule absorption by PDMS in the context of drug response bioassays. Biochemical and Biophysical Research Communications 482, 323–328 (2017).

17. Toepke, M. W. & Beebe, D. J. PDMS absorption of small molecules and consequences in microfluidic applications. Lab Chip 6, 1484–1486 (2006).

18. Mukhopadhyay, R. When PDMS isn’t the best. Anal. Chem. 79, 3248–3253 (2007).

19. Li, Y. & Kumacheva, E. Hydrogel microenvironments for cancer spheroid growth and drug screening. Sci. Adv. 4, eaas8998 (2018).

20. Lee, J. M. et al. Generation of uniform-sized multicellular tumor spheroids using hydrogel microwells for advanced drug screening. Sci Rep 8, 17145 (2018).

21. Shen, C., Li, Y., Wang, Y. & Meng, Q. Non-swelling hydrogel-based microfluidic chips. Lab Chip 19, 3962–3973 (2019).

22. Humayun, M. et al. Elucidating cancer-vascular paracrine signaling using a human organotypic breast cancer cell extravasation model. Biomaterials 270, 120640 (2021).

23. Haessler, U., Kalinin, Y., Swartz, M. A. & Wu, M. An agarose-based microfluidic platform with a gradient buffer for 3D chemotaxis studies. Biomed Microdevices 11, 827–835 (2009).

24. Liu, T. et al. A microfluidic device for characterizing the invasion of cancer cells in 3-D matrix. Electrophoresis 30, 4285–4291 (2009).

25. Pluen, A., Netti, P. A., Jain, R. K. & Berk, D. A. Diffusion of Macromolecules in Agarose Gels: Comparison of Linear and Globular Configurations. Biophysical Journal 77, 542–552 (1999).

26. Zarrintaj, P. et al. Agarose-based biomaterials for tissue engineering. Carbohydrate Polymers 187, 66–84 (2018).

27. Reinfeld, B. I. et al. Cell-programmed nutrient partitioning in the tumour microenvironment. Nature 593, 282–288 (2021).

28. Sabatini, D. A. Sorption and Intraparticle Diffusion of Fluorescent Dyes with Consolidated Aquifer Media. Groundwater 38, 651–656 (2000).

29. Gustafsson, J. et al. Metabolic collaboration between cells in the tumor microenvironment has a negligible effect on tumor growth. The Innovation 5, 100583 (2024).

30. Verhagen, N. et al. Comparison of l-tyrosine containing dipeptides reveals maximum ATP availability for l-prolyl-l-tyrosine in CHO cells. Engineering in Life Sciences 20, 384–394 (2020).

31. Prunet, A. et al. A new agarose-based microsystem to investigate cell response to prolonged confinement. Lab Chip 20, 4016–4030 (2020).

32. Bhatia, S. N. & Ingber, D. E. Microfluidic organs-on-chips. Nat Biotechnol 32, 760–772 (2014).

33. Tomasi, R. F.-X., Sart, S., Champetier, T. & Baroud, C. N. Individual Control and Quantification of 3D Spheroids in a High-Density Microfluidic Droplet Array. Cell Reports 31, 107670 (2020).

34. Parihar, P. et al. An Overview of Advancements and Technologies in Vascularization Strategies for Tumor-On-A-Chip Models. Advanced therapeutics (2024).

35. Joshi, I. M. et al. Microengineering 3D Collagen Matrices with Tumor-Mimetic Gradients in Fiber Alignment. Advanced Functional Materials 34, 2308071 (2024).

36. He, J. et al. Gelatin Methacryloyl Hydrogel, from Standardization, Performance, to Biomedical Application. Advanced Healthcare Materials 12, 2300395 (2023).

37. Piironen, K., Haapala, M., Talman, V., Järvinen, P. & Sikanen, T. Cell adhesion and proliferation on common 3D printing materials used in stereolithography of microfluidic devices. Lab Chip 20, 2372–2382 (2020).

38. Hart, C., Didier, C. M., Sommerhage, F. & Rajaraman, S. Biocompatibility of Blank, Post-Processed and Coated 3D Printed Resin Structures with Electrogenic Cells. Biosensors 10, 152 (2020).

39. Rios, A. C. et al. Intraclonal Plasticity in Mammary Tumors Revealed through Large-Scale Single-Cell Resolution 3D Imaging. Cancer Cell 35, 618–632.e6 (2019).

40. Zhang, J. et al. 13C Isotope-Assisted Methods for Quantifying Glutamine Metabolism in Cancer Cells. in Methods in Enzymology vol. 542 369–389 (Elsevier, 2014).

