## Supplementary_data for "Integrating 3D Tumor Models and Microfluidics for Precise Metabolic Control"

[1] Univ Lyon, Univ Claude Bernard Lyon 1, CNRS, Institut Lumière Matière, F-69622, VILLEURBANNE, France.
[2] µFab Plateforme microfabrication, Institut Convergence Plascan, Centre de Recherche en Cancérologie de Lyon, INSERM U1052-CNRS UMR5286, Centre Léon Bérard, Université de Lyon, Université Claude Bernard Lyon1, 69008 Lyon, France

Keywords: tumor-on-chip, hydrogel microenvironement, L-glutamine metabolism, colon cancer

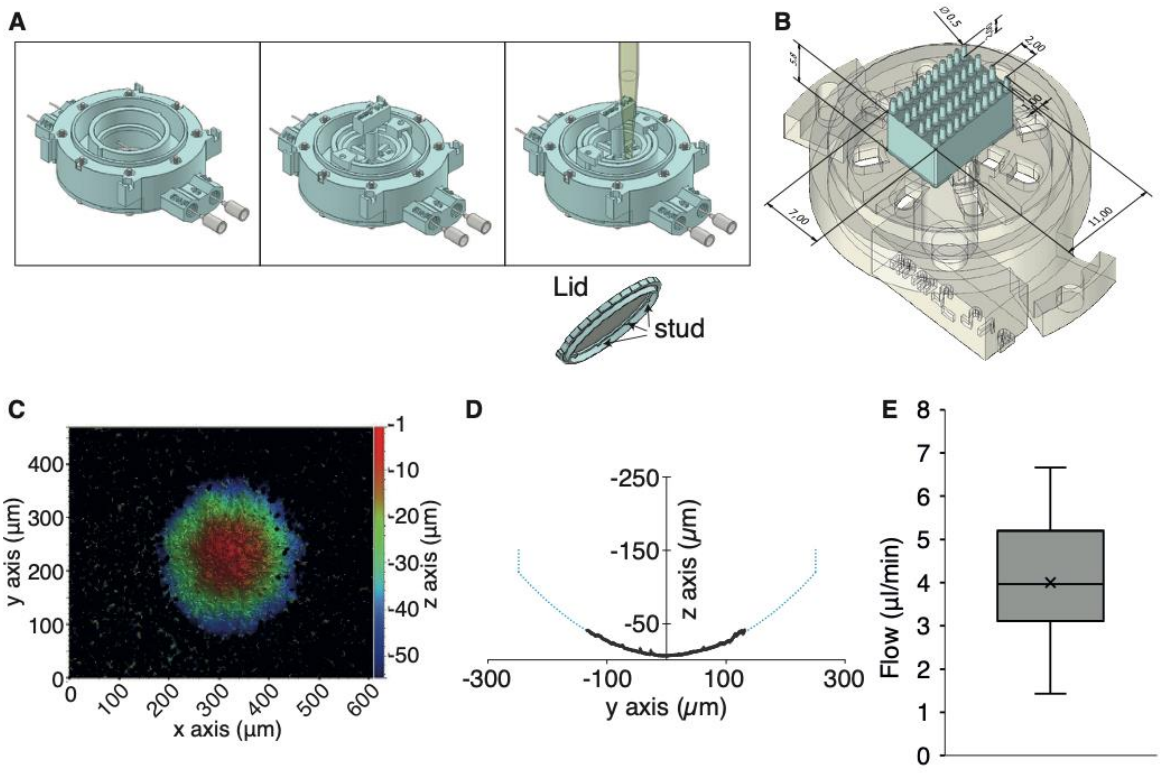


**Figure S1. MilliFlow assembly, microwell geometry and flow rate characterization.** (**A**) Schematic sequence illustrating the assembly of the microneedles and the stamp within the chip and the agarose molding (from left to right). The lid illustration shows supporting studs resting on the central part of the chip. The spacing between adjacent studs creates a 1-mm-high free space between the lid and the chip base, enabling efficient gas exchange essential for cell culture. (**B**) Technical drawing of the stamp showing the dimensions of the rectangular seeding chamber mold and the microneedles, as well as their arrangement in an array. (**C**) Representative optical profilometry image (Contour GT, Bruker; objective G = 10×) of a printed microwell tip (500 µm in diameter). (**D**) Mean ± SD of the measured curvature profile along the *y*-axis (black solid line) and corresponding extrapolated fit (blue dotted line), confirming the diameter of 500 um ($n$ = 11 microwells, $N$ = 1 experiment). (**E**) Time-dependent flow rate characterization by hourly weight measurement of the collected medium. Data are presented as mean ± SD, $N$ = 3 independent experiments.

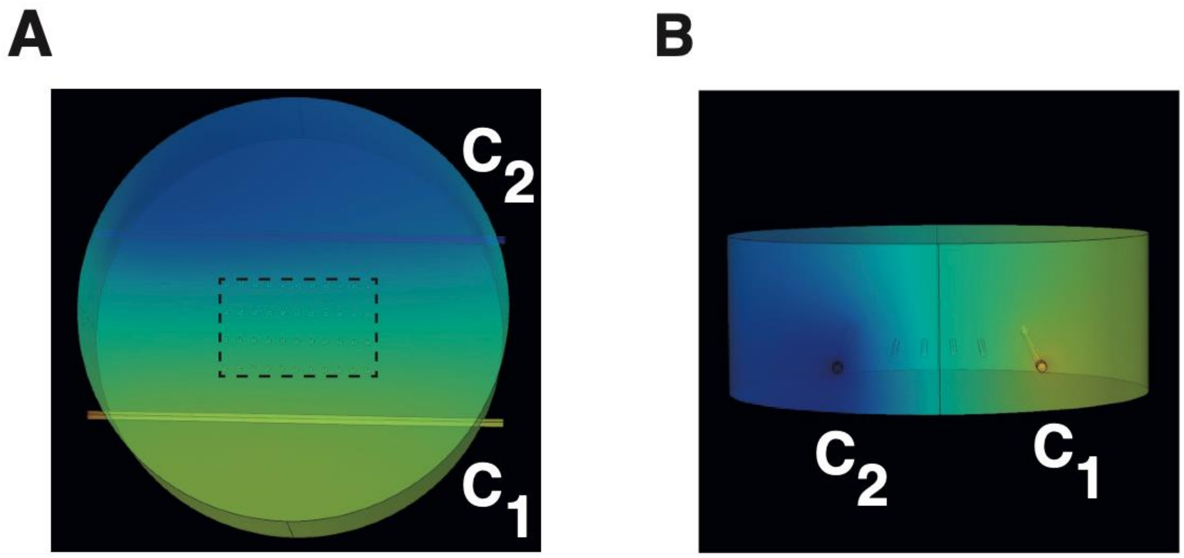


**Figure S2. Homogeneous gradient across the seeding chamber surface.** Three-dimensional COMSOL simulations showing the SRB concentration gradient established 24 h after perfusion initiation. Two conditions are shown: C1, perfusion with SRB-containing medium, and C2, perfusion with medium alone. (**A**) Top view and (**B**) transverse view of the simulated domain. The rectangle indicates the location of the culture (seeding) chamber, while circles mark the positions of the microwells. Simulations were performed using COMSOL Multiphysics, incorporating the geometrical parameters of the device, hydrogel properties, and SRB transport characteristics. The color map ranges from yellow (high SRB concentration) to blue (low SRB concentration).

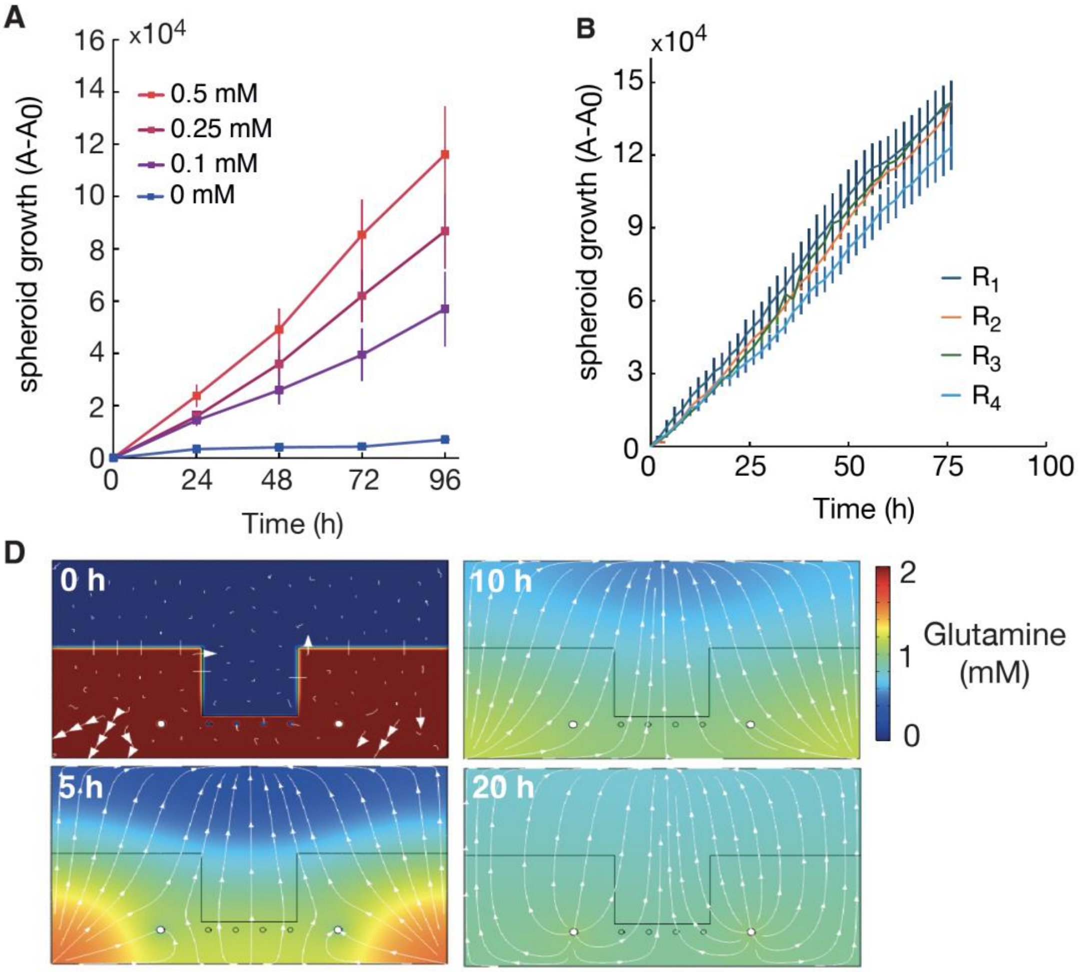


**Figure S3. Control experiments validating spheroid growth behavior under defined L-glutamine conditions.** (A) Growth curves of HCT116 spheroids cultured in ultra–low attachment (ULA) plates in medium containing 0, 0.05, 0.1, 0.25, or 0.5 mM L-glutamine. Mean +/- SEM, N=3 (B) Growth curves of HCT116 spheroids cultured in the MilliFlow3D chip perfused with medium supplemented with 1 mM L-glutamine in both channels, showing no significant growth differences between rows in the absence of a gradient. Mean +/- SEM, N=1 (C) 2D diffusion-simulated cross-sectional views illustrating L-glutamine distribution over time within the agarose gel containing the seeding chamber, when both perfusion channels are supplied with 1 mM L-glutamine. The color map indicates L-glutamine concentration and arrows represent diffusion fluxes.

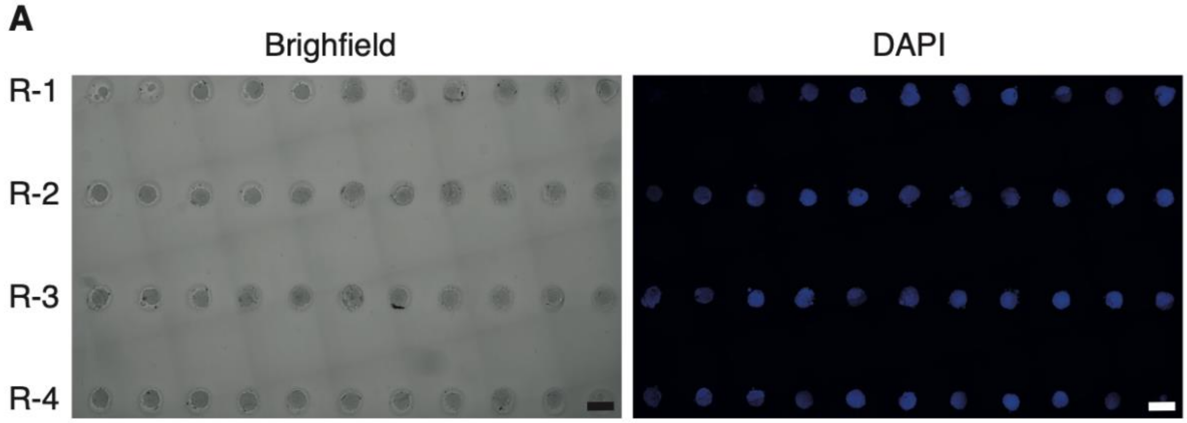


**Figure S4.** (A) Stitched brightfield image of the seeding chamber and corresponding confocal image of spheroids stained with DAPI to visualize nuclei. Scale bar: 500 µm
